# Collective AI use is associated with researcher engagement: Real-time evidence from a scientific conference

**DOI:** 10.64898/2026.01.26.701639

**Authors:** Hiroyuki Okada, Shigeto Seno, Ung-il Chung, Naganari Okura

**Affiliations:** Department of Clinical Biotechnology, Center for Disease Biology and Integrative Medicine, Graduate School of Medicine, the University of Tokyo, Tokyo, Japan; Department of Bioinformatic Engineering, Graduate School of Information Science and Technology, the University of Osaka, Osaka, Japan; Department of Bioengineering, Graduate School of Engineering, the University of Tokyo, Tokyo, Japan; Department of Frontier Research in Tumor Immunology, Graduate School of Medicine, the University of Osaka, Osaka, Japan

## Abstract

Recent large-scale bibliometric analyses suggest that individual AI use can increase productivity while reducing downstream engagement and topic diversity. Here we ask whether collective AI deployment is associated with shared engagement. Using an Audience Response Engagement (ARE) system at NGS Expo 2025 (N=110 biomedical researchers), we captured real-time consensus and generated updated visualizations within minutes. Our data reveal a substantial gap between adoption and transparency: 93.6% of researchers use AI at least weekly, yet only 5.5% consistently disclose this usage—a 17-fold disparity. This pattern is consistent with systemic policy uncertainty (39.1% report unclear guidelines). Behavioral clustering identifies a “High-Concern” group (31%) as a candidate for targeted interventions: highest productivity yet lowest disclosure. These findings suggest that collective AI deployment in physical settings is associated with shared engagement.

---

Recent work suggests that individual AI use increases productivity while reducing the diversity of scientific inquiry^1^. We asked whether collective AI deployment—real-time, shared use in physical settings—might support rather than diminish researcher engagement. Here we examine this question at a scientific conference.

In-person scientific conferences offer a natural testing ground. Post-pandemic, these gatherings face pressure to justify travel costs when virtual alternatives exist^2^,^3^. Current formats—static presentations followed by brief Q&A—provide limited engagement□. Yet physical co-presence enhances shared excitement, spontaneous interaction, and collective discovery in real-time more than virtual meetings.

We hypothesized that AI could amplify this energy. Rather than isolating researchers into productivity silos, collective AI deployment could engage audiences as active collaborators— generating real-time consensus, co-created visualizations, and immediate research outputs.

Compounding the isolation problem is a transparency gap in AI use. Generative AI tools (ChatGPT, Claude, Gemini) have permeated research workflows since late 2022□,□; yet institutional policies for disclosure of AI usage lag behind adoption□. Publisher requirements vary: Nature mandates Methods section disclosure□; Elsevier requires acknowledgment□; COPE has issued guidance^1^□, but implementation remains inconsistent. This policy vacuum creates uncertainty about appropriate disclosure practices.

Empirical data on AI practices among researchers remains sparse. Existing studies rely on online surveys with self-selected samples□,□ or focus on narrow domains. No studies have captured real-time, in-person consensus on AI practices from active researchers at a scientific meeting.

Here we contribute (i) a live, reproducible pipeline for real-time audience-in-the-loop analysis, and (ii) quantitative estimates of the adoption–disclosure gap with behavioral profiling in an in-person setting.

We deployed an Audience Response Engagement (ARE) system at NGS Expo 2025 (October 29–30, Nara, Japan), a next-generation sequencing conference (Fig. 1a). The system uses widely accessible, low-barrier tools (Google Forms, Google Sheets, Python, Marp) to enable live survey, analysis, and slide updates during the presentation.

**Figure 1.**
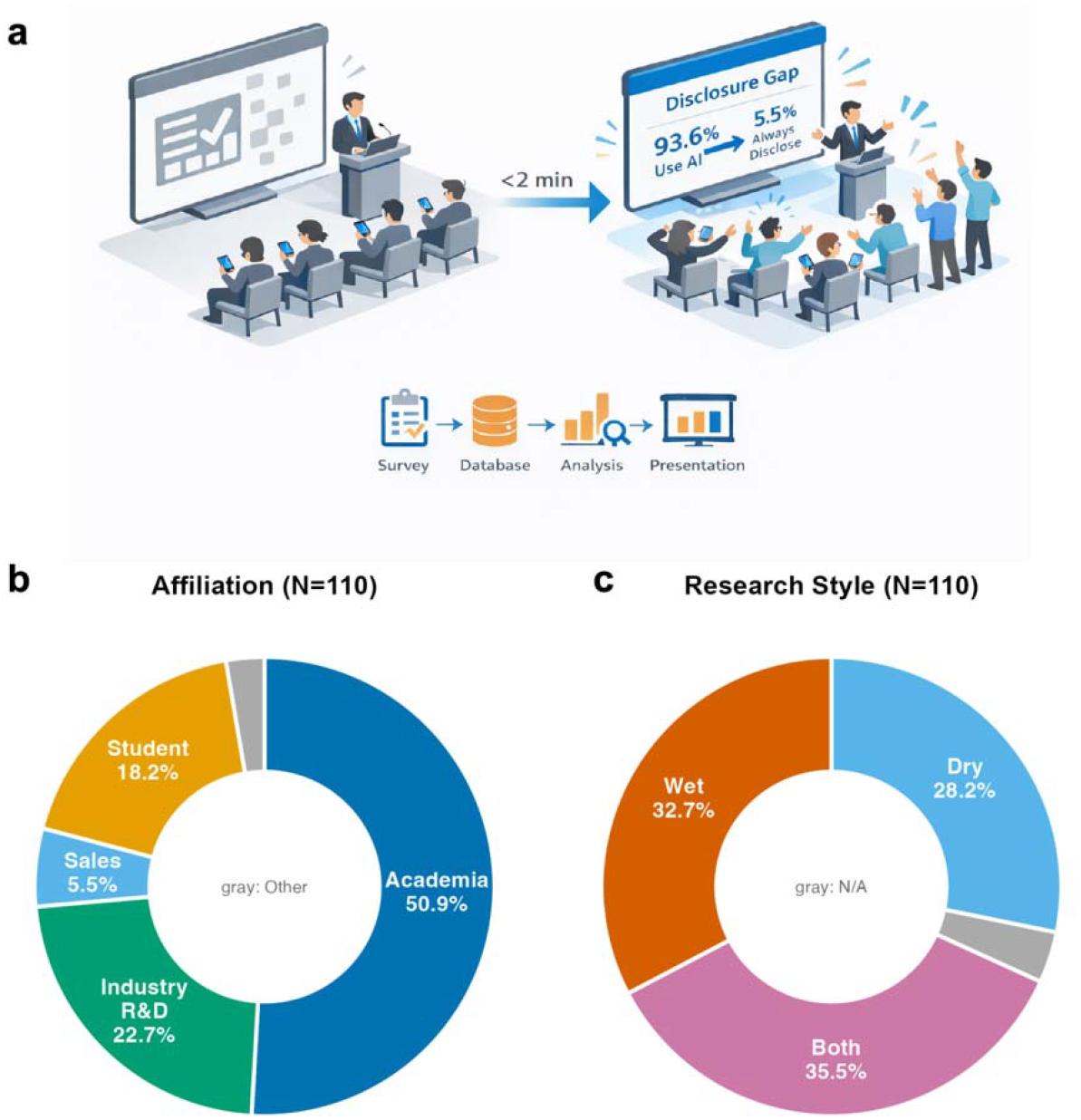
ARE System Overview and Participant Characteristics. (a) System architecture: QR code distribution to audience, data flow through Google Forms and Sheets, Python analysis, and live Marp slide updates. (b) Participant affiliation distribution (N=110). (c) Research style distribution.

Our 13-question bilingual survey (Japanese/English; Extended Data 1) assessed participant demographics, institutional context (AI guidelines), AI usage patterns (frequency, tools, use cases), and disclosure practices. Question 10 used a 7-point equal-interval scale for productivity assessment, enabling parametric statistical analysis.

Participants took part voluntarily and anonymously. We obtained publication consent via an opt-out item (Q13). We received 116 responses; after excluding 3 test entries and 3 who declined publication, 110 remained for analysis. Participants were predominantly from academia (50.9%) and industry R&D (22.7%), with students comprising 18.2% (Fig. 1b). Research styles were balanced across experimental (32.7%), computational (27.3%), and combined approaches (35.5%; Fig. 1c).

Real-time engagement demonstrates technical feasibility. The ARE system operated without interruptions throughout the presentation. The first response arrived 47 s after QR code display; response rate peaked at 23 submissions within a 3-min window. Of valid responses, 60.2% (n=68) arrived during the 15-min talk, with 39.8% (n=45) arriving in the subsequent 2-hour window. Five figures were updated live with latency under 2 min from data collection to display.

Participants observed their collective input emerge in real-time—responses visualized, concerns aggregated, consensus formed. This shared experience of co-creating research output exemplifies a form of collective engagement that may be difficult to reproduce in fully virtual formats.

High AI adoption with consistent productivity gains. AI adoption was widespread: 73.6% reported daily use and 20.0% weekly use, meaning over 93% use AI tools at least weekly. ChatGPT dominated (92.0%), with substantial adoption of Gemini (39.8%) and Claude (21.2%; Extended Data 1). Multiple tool use was common, indicating active exploration of the AI ecosystem.

Productivity gains were consistent (Fig. 2a). On our 7-point equal-interval scale (1=significantly decreased to 7=significantly increased, 4=neutral), respondents reported a mean of 6.17±0.78 (N=108). This represents Cohen’s d=2.78, significantly above the neutral midpoint. Of respondents, 97.2% reported positive productivity impact (scores 5–7), with 82% reporting “moderately” or “significantly” increased productivity (scores 6–7). No respondents reported decreased productivity. Higher usage frequency predicted greater gains (Fig. 2b).

**Figure 2.**
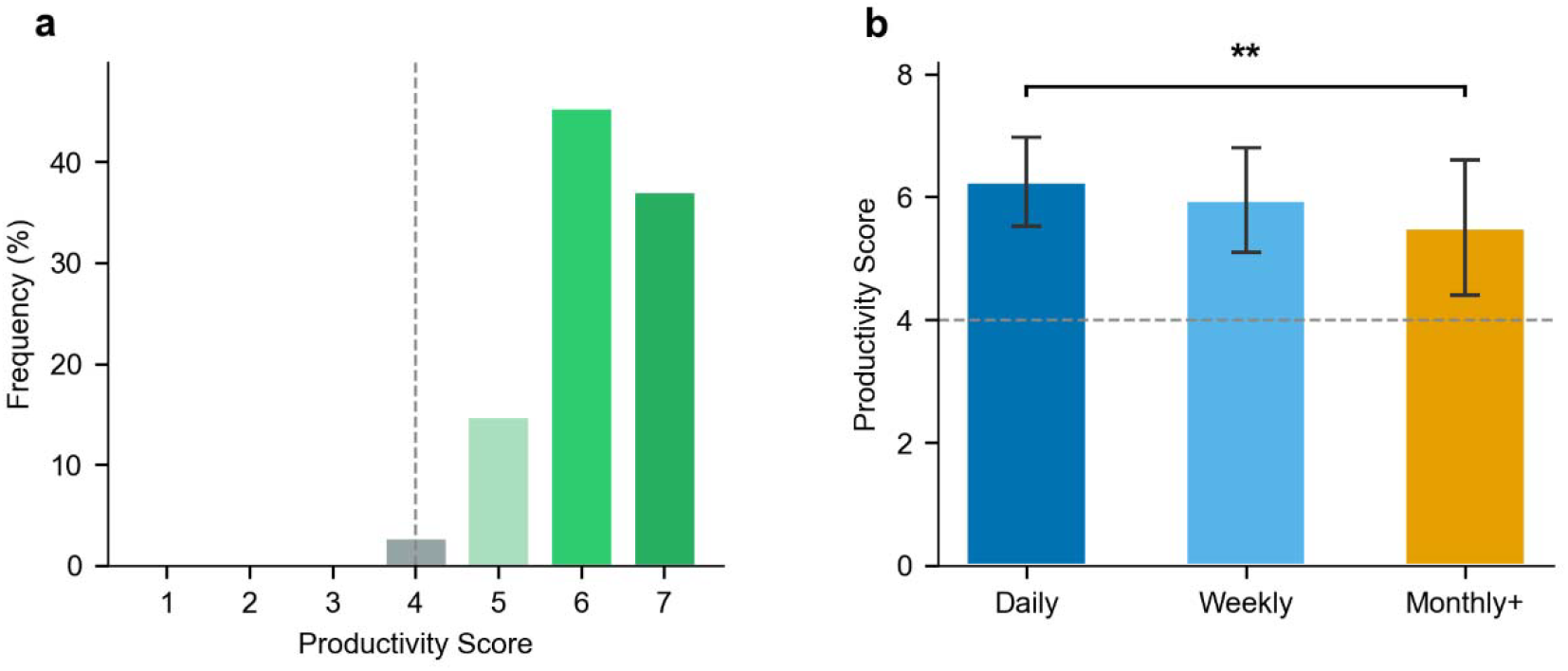
AI Productivity Impact. (a) Self-reported productivity impact on 7-point equal-interval scale (1=significantly decreased, 4=neutral, 7=significantly increased; N=108). Dashed line indicates neutral. Mean=6.17±0.78, Cohen’s d=2.78. (b) Mean productivity score (±SD) by AI usage frequency. **p<0.01; bracket indicates Daily vs Monthly+ comparison.

The 17-fold disclosure gap. A substantial disparity emerged between AI adoption and transparency (Fig. 3a). While 93.6% used AI at least weekly, only 5.5% consistently disclosed this usage in publications (Q11=“Always disclose”). This 17-fold gap warrants attention as AI’s role in research faces increasing scrutiny.

**Figure 3.**
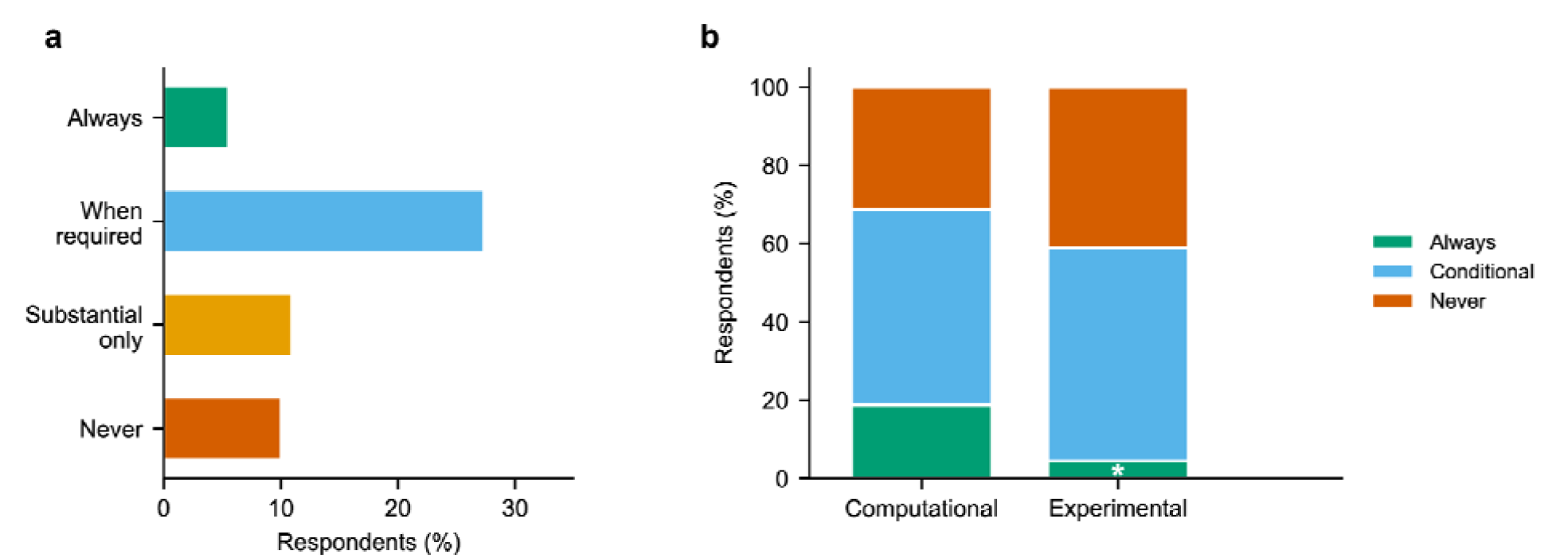
AI Disclosure Practices. (a) Distribution of disclosure practices among respondents with publication experience (N=59). (b) Disclosure patterns by research style (Computational vs Experimental; N=60). Computational researchers showed higher “Always disclose” rates (18.8% vs 4.5%, Cohen’s h=0.47, 96% posterior probability).

This pattern is consistent with systemic policy uncertainty (Extended Data 2). When asked about institutional AI guidelines, 39.1% reported unclear or absent policies. An additional 22.7% were unaware whether guidelines existed. Only 25.5% reported clear guidance. When policies are ambiguous, researchers may default to non-disclosure.

Disclosure patterns varied by research style (Fig. 3b). Among respondents with publication experience (N=60), computational researchers (“Dry”) disclosed AI use at substantially higher rates than experimental researchers: 18.8% vs 4.5% (Cohen’s h=0.47, medium effect; Fisher’s exact p=0.112). Bayesian analysis using Beta-Binomial conjugate priors yielded a 96% posterior probability that computational researchers disclose at higher rates than others. This suggests that transparency norms cultivated in computational communities—where software acknowledgment and code sharing are standard practice^11^,^12^—may transfer to AI disclosure practices.

Behavioral clustering reveals distinct user profiles. To characterize behavioral heterogeneity, we performed K-means clustering (k=4, selected by silhouette analysis; see Methods) on standardized behavioral features (Fig. 4a). Four clusters emerged, defined by usage patterns rather than demographics.

**Figure 4.**
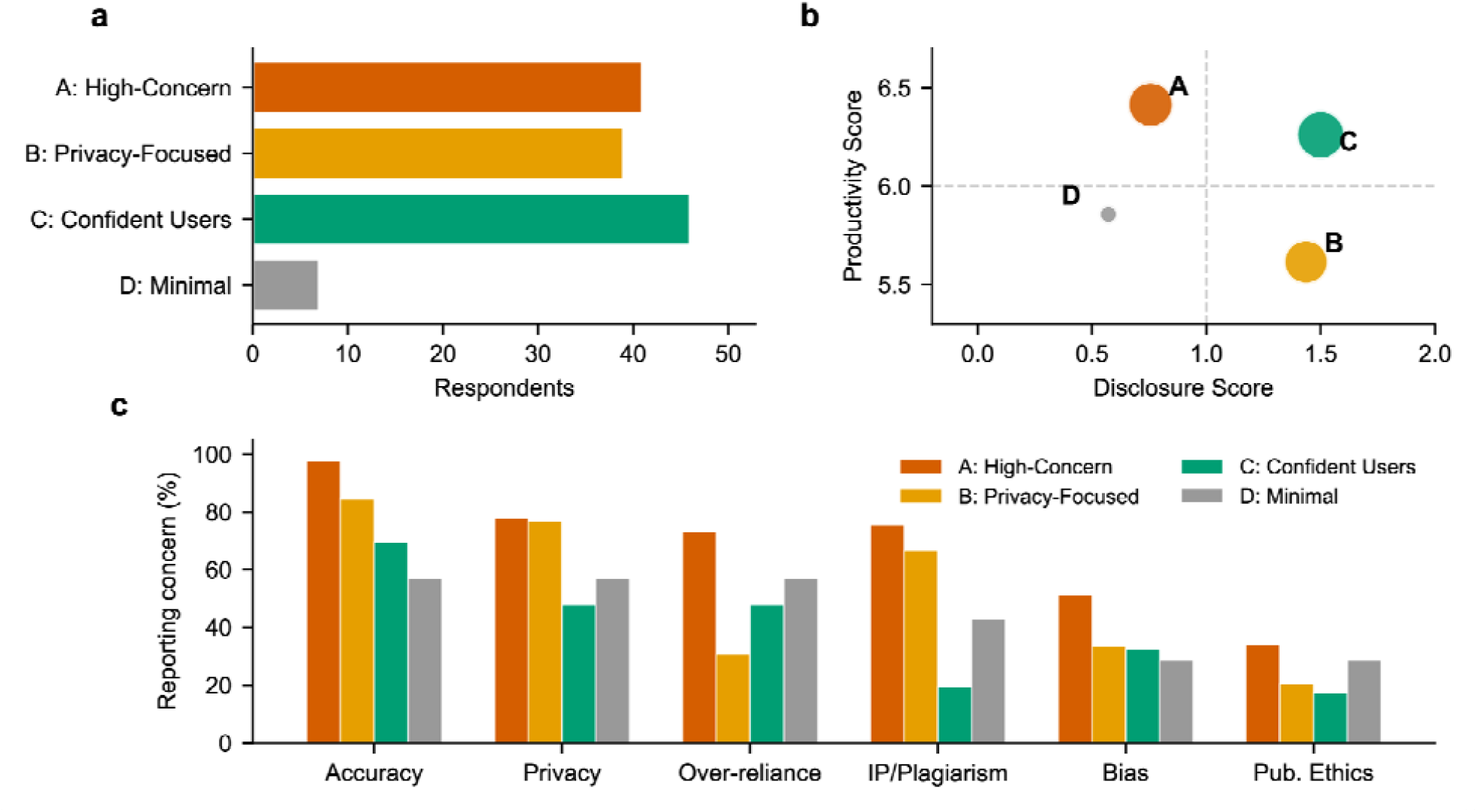
Behavioral Clustering Analysis. (a) Four clusters identified by K-means clustering (k=4). (b) Cluster positions in productivity-disclosure space; bubble size proportional to cluster size. (c) Concern profiles by cluster.

The “High-Concern” cluster (Cluster A, 31%, n=34) showed the largest gap between productivity and disclosure: highest productivity scores (mean 6.41) combined with lowest disclosure rates, and elevated concerns across all categories including accuracy, privacy, intellectual property, and over-reliance (Fig. 4b,c). This group may represent a candidate for targeted interventions.

Conversely, “Confident Users” (Cluster C, 35%, n=38) achieved comparable productivity (mean 6.28) with the highest disclosure rates and fewest concerns. This group used more diverse AI tools and may serve as transparency role models. The “Privacy-Focused” cluster (Cluster B, 29%, n=32) showed moderate engagement with specifically elevated privacy concerns. The “Minimal” cluster (Cluster D, 5%, n=6) reported limited AI engagement overall. These distinct profiles suggest that differentiated approaches—rather than blanket policies—might be more effective for addressing disclosure gaps.

## Discussion

These findings illustrate a deployment mode consistent with increased shared engagement. Hao et al.^1^ documented a trade-off: individual AI use boosts productivity but contracts collective diversity. The ARE system suggests an alternative deployment mode—collective use in physical settings—that may preserve productivity gains while fostering shared engagement rather than isolation.

Rather than using AI for individual acceleration, we used it to amplify a physical gathering. The 110 researchers became co-creators of research output, observing consensus emerge in real-time. Whether this approach can mitigate the isolation documented elsewhere^1^ requires longitudinal study with network measures.

The 17-fold disclosure gap highlights a need for clearer institutional policies. Our data suggest this gap is consistent with policy ambiguity. When 93.6% of peers use AI regularly, disclosure may warrant normalization.

This study has limitations. Single-event sampling (genomics, Japan) limits generalizability. Self-selection and self-report biases may affect results, though anonymity mitigates concerns. The disclosure subgroup analysis (N=60) was underpowered (∼36% power); N≈200 per group needed for 80% power. Validation across diverse conferences is needed.

This approach uses widely accessible tools. Google Forms, Python, and Marp enable implementation with minimal technical barriers.

Hao et al.^1^ showed that individual AI use expands productivity but contracts scientific diversity. Our findings suggest that deployment mode matters: collective AI use in physical settings is associated with shared engagement, where researchers become co-creators rather than isolated users. Future work should examine whether such collective deployment can preserve productivity gains while mitigating the diversity contraction observed in individual use.

## Methods

### Study design

We conducted a cross-sectional survey at NGS Expo 2025 (October 29–30, Nara, Japan). Participation was voluntary and anonymous. This study did not require institutional ethics review under the University of Tokyo’s guidelines, as it constituted minimal-risk research: an anonymous survey of researchers at a professional conference, with voluntary participation and an explicit opt-out option for publication inclusion. Participants were informed of the study purpose via the survey introduction; consent was documented through voluntary completion.

### Survey instrument

We developed a 13-question bilingual survey (Japanese/English) covering demographics, institutional context, AI usage patterns, practices, concerns, and publication consent (Table 1). Question 10 assessed productivity using a 7-point equal-interval scale (1=significantly decreased, 4=neutral, 7=significantly increased), enabling parametric analysis. Full questionnaire in Table S1.

**Table 1.** Survey Question Summary. The 13-item bilingual questionnaire (Japanese/English) administered at NGS Expo 2025 (N=110). Question 10 uses a 7-point equal-interval scale enabling parametric statistical analysis. Multiple choice questions (Q5, Q7, Q8) allowed multiple selections.

| # | Domain | Question Topic | Format |
| --- | --- | --- | --- |
| 1 | Demographics | Primary affiliation | Single choice |
| 2 | Demographics | Research style (Wet/Dry/Both) | Single choice |
| 3 | Context | Institutional AI guidelines | Single choice |
| 4 | Usage | AI usage frequency | Single choice |
| 5 | Usage | AI tools used | Multiple choice |
| 6 | Usage | AI cost funding source | Single choice |
| 7 | Usage | Primary use cases | Multiple choice |
| 8 | Concerns | Main concerns about AI | Multiple choice |
| 9 | Practices | AI output verification level | Single choice |
| 10 | Practices | Productivity impact (1-7 scale) | Equal-interval |
| 11 | Practices | AI disclosure in publications | Single choice |
| 12 | Practices | AI adoption timeline | Single choice |
| 13 | Consent | Publication consent | Opt-out |

### ARE system architecture

The system pipeline was: Google Forms → Google Sheets → Sheets API v4 → Python (gspread, pandas, matplotlib) → Marp CLI. The script polled for new responses at 30-second intervals. Mean latency from response submission to slide update was under 2 minutes. Five figures were updated in real-time during the presentation. This pipeline uses only free-tier cloud services (Google Forms, Google Sheets) and open-source software (Python 3.8+, Marp CLI), enabling replication without licensing costs. No proprietary software or paid subscriptions are required.

### Statistical analysis

Primary outcomes were disclosure rate and productivity score. Two participants did not respond to the productivity question; analyses used complete cases (N=108 for productivity, N=110 for other variables). We treated the 7-point productivity scale as approximately interval; results were consistent under non-parametric testing (Wilcoxon signed-rank test confirmed median significantly above neutral). We calculated descriptive statistics for categorical (frequencies) and continuous (means±SD) variables. We tested frequency– productivity associations using one-way ANOVA with post-hoc Tukey HSD, and disclosure associations using chi-square tests and Fisher’s exact test for 2×2 tables. Effect sizes: Cohen’s d (continuous), Cohen’s h (proportions). We used Bayesian inference for disclosure comparison with Beta-Binomial conjugate priors (uniform Beta(1,1) priors) and Monte Carlo sampling (n=100,000). For behavioral clustering, we applied K-means (k=4) to z-scored features (productivity, disclosure, concern scores, tool diversity); k was selected by silhouette analysis (k=3–5 tested; k=4 yielded highest score=0.31); results were robust to random initialization (n_init=10, random_state=42). Significance threshold α=0.05 (two-tailed). Software: Python 3.8+ (scipy, scikit-learn).

## Data availability

Anonymized survey responses (N=110) will be provided when this article is published.

## Acknowledgements

We gratefully acknowledge the NGS Expo 2025 organizing committee. We thank the 110 conference participants who contributed their perspectives.

## Funding

This work was supported by JSPS KAKENHI (22KK0272, 22H05084, 23K15736, 25K12430); DxPoly Program (JPMXP1122714694); ERATO SAKAI Real and Abstract Gels Project (JPMJER2401); Takeda Science Foundation (2024); Mochida Memorial Foundation for Medical and Pharmaceutical Research (2022); and Astellas Foundation for Research on Metabolic Disorders (2023) to H.O.

## Author Contributions

H.O.: Conceptualization, Methodology, Software, Formal Analysis, Investigation, Writing— Original Draft, Visualization, Funding acquisition. S.S.: Conceptualization, Methodology, Writing—Review & Editing. U.C.: Writing—Review & Editing. N.O.: Conceptualization, Writing—Review & Editing.

## Competing Interests

None declared.

## AI Disclosure

Claude Opus 4.5 (Anthropic) assisted with literature search, outline structuring, and statistical interpretation. All analyses were independently verified by H.O. No AI-generated text was used without critical review. No generative AI tools were used to create or modify figures or images except Figure 1a. Data visualizations (Figures 1b,c,2–4) were generated using Python (Matplotlib/Seaborn).

**Extended Data 1.**
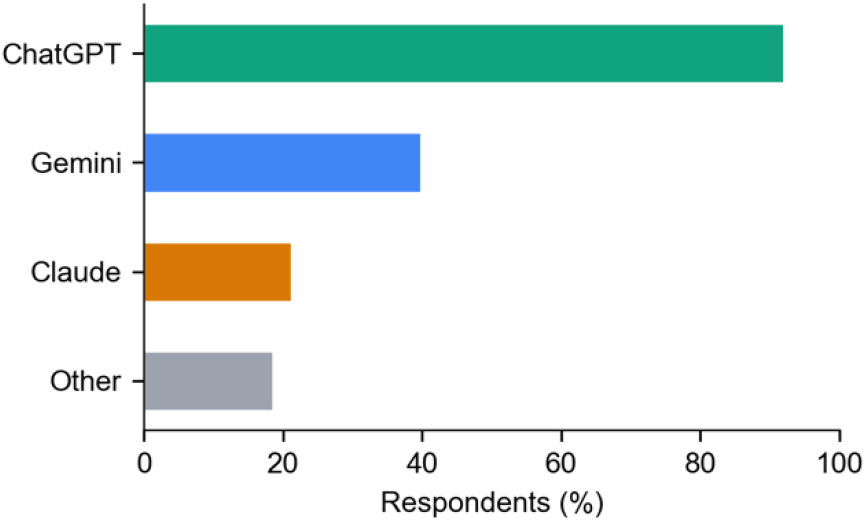
AI Tools Distribution. AI tools used by participants (N=110; multiple selection allowed).

**Extended Data 2.**
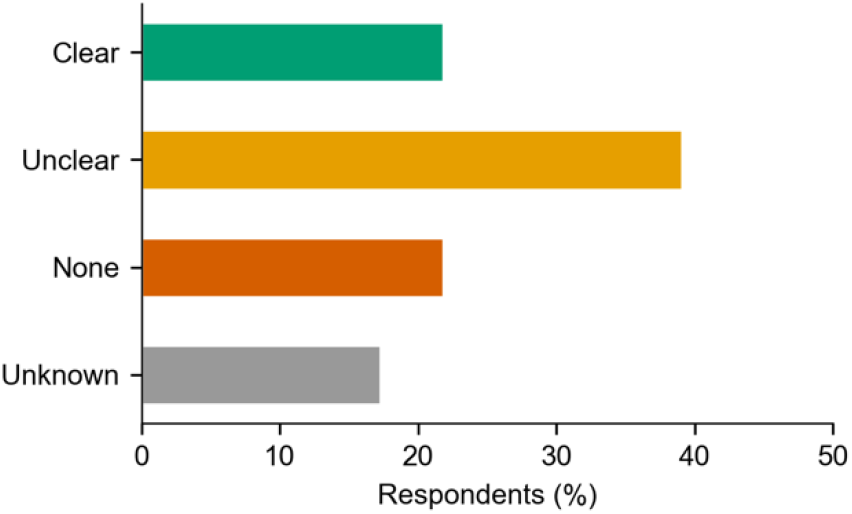
Institutional AI Guidelines. Institutional AI guideline status as reported by participants (N=110).

## Notes

### Competing Interest Statement

The authors have declared no competing interest.

## References

1. Hao, Q. et al. Artificial intelligence tools expand scientists’ impact but contract science’s focus. Nature 10.1038/s41586-025-09922-y (2026).

2. Remmel, A. Scientists want virtual meetings to stay after the COVID pandemic. Nature 591, 185–186 (2021).

3. Skiles, M. et al. Conference demographics and footprint changed by virtual platforms. Nat. Sustain. 5, 149–156 (2022).

4. Jerardi, K. E. et al. Evaluating the impact of interactive educational conferences. Perspect. Med. Educ. 2, 349–355 (2013).

5. Owens, B. How Nature readers are using ChatGPT. Nature 615, 20 (2023).

6. Van Noorden, R. & Perkel, J. M. AI and science: what 1,600 researchers think. Nature 621, 672–675 (2023).

7. Rasky, E. Generative AI Policy in Higher Education: A Preliminary Survey. CIGI Report (2023).

8. Nature Editorial. Tools such as ChatGPT threaten transparent science. Nature 613, 612 (2023).

9. Elsevier. The use of AI and AI-assisted technologies in scientific writing. Elsevier Policy (2023).

10. COPE Council. Authorship and AI tools: COPE Position Statement (2023).

11. Cadwallader, L. et al. Advancing code sharing in the computational biology community. PLoS Comput. Biol. 18, e1010193 (2022).

12. Ince, D. C. et al. The case for open computer programs. Nature 482, 485–488 (2012).

